# Reconstitution of Integral Membrane Proteins at Catenoid Necks via the Synthetic Membrane Transfer

**DOI:** 10.1101/2025.08.23.671895

**Authors:** Hemraj Meena, Rafal Zdanowicz, Tobias Bock-Bierbaum, Till Stephan, Sofie Dorothea Ottsen Knudsen, Stefan Jakobs, Oliver Daumke, Per Amstrup Pedersen, Nicola De Franceschi

**Affiliations:** IMol Polish Academy of Sciences, Warsaw, Poland; Structural Biology, Max Delbrück Center for Molecular Medicine in the Helmholtz Association (MDC), Berlin, Germany; Research Group Mitochondrial Structure and Dynamics, Max Planck Institute for Multidisciplinary Sciences, Göttingen, Germany; High Resolution Microscopy, Clinic for Neurology, University Medical Center Göttingen, Göttingen, Germany; Buchmann Institute for Molecular Life Sciences, Goethe University Frankfurt am Main, Germany; Institute of Molecular Biosciences, Goethe University Frankfurt am Main, Germany; Department of Biology, University of Copenhagen, Copenhagen, Denmark; Translational Neuroinflammation and Automated Microscopy, Fraunhofer Institute for Translational Medicine and Pharmacology ITMP, Göttingen, Germany

**Keywords:** Integral membrane protein, reconstitution, GUV, aquaporin, Mic10, dumbbells

## Abstract

Functional reconstitution of integral membrane proteins (IMPs) in giant unilamellar vesicles (GUVs) remains a major bottleneck in bottom-up synthetic biology, particularly when combined with controlled membrane geometry. Here, we introduce the Synthetic Membrane Transfer (SMT), a detergent-free strategy that exploits exchange from styrene–maleic acid nanoparticles to enable direct insertion of IMPs into GUV membranes during vesicle formation. SMT operates under mild conditions and is compatible with inverted emulsion protocols, allowing simultaneous lumenal encapsulation and membrane reshaping. Here, using the SMT we demonstrate functional reconstitution of two structurally distinct IMPs, aquaporin-12 and the mitochondrial protein Mic10 in GUVs that are simultaneously deformed into dumbbell geometries featuring catenoid membrane necks. Mic10 forms oligomeric clusters and exhibits robust enrichment at neck regions, providing direct evidence of curvature-dependent sorting of a transmembrane hairpin protein in a minimal system. Together, these results establish SMT as a versatile platform for spatially controlled assembly of membrane protein modules and enable interrogation of transmembrane protein behavior in complex membrane geometries, providing a foundation for bottom-up reconstitution of multiprotein assemblies at catenoid membrane necks.

## Introduction

Integral membrane proteins (IMPs) are essential components of biological membranes, where they perform crucial functions including energy generation (Salmonowicz, 2025), transport (Alam, 2023), signaling (Ripoll, 2025) and the organization of multiprotein assemblies (Nakamura, 2024). They are also highly represented among the targets of clinically used drugs (Santos, 2017). Reconstituting IMPs in simplified membrane systems is therefore a central goal of bottom-up synthetic biology and membrane biophysics.

Giant unilamellar vesicles (GUVs) provide a particularly powerful platform for studying membrane proteins, owing to their cell-like dimensions, optical accessibility, and compatibility with real-time fluorescence microscopy (Litschel, 2021; Fragasso, 2021). GUVs are also widely used as chassis for the construction of synthetic cells (Olivi, 2021; Li, 2025). While generating protein-free GUVs from phospholipids is straightforward using methods such as electroformation (Streicher, 2009; Varnier, 2010; Aimon, 2011; Gaul, 2015; Almendro-Vedia, 2017, Kumar, 2024), gel-assisted swelling (Souissi, 2021; Tosaka, 2024; Cooper, 2025), or inverted emulsion (Yanagisawa, 2011; Altamura, 2017; Suzuki, 2023), the functional reconstitution of transmembrane proteins into GUVs remains a major bottleneck. Existing approaches typically require extensive case-by-case optimization, suffer from poor encapsulation efficiency, or rely on non-physiological membrane compositions (Biner, 2016, Stępień, 2023). These limitations become particularly restrictive when reconstitution must be combined with controlled membrane geometry.

Nanodisc-based strategies have emerged as powerful tools for stabilizing and characterizing IMPs in defined lipid environments (Sligar, 2021). In parallel, synthetic polymer–lipid nanoparticles offer attractive alternatives to MSP-based nanodiscs. Styrene–maleic acid (SMA) is an amphipathic copolymer composed of hydrophobic styrene and hydrophilic maleic acid groups, whose relative abundance and distribution influence its solubility, conformation, and membrane-interacting properties (Dörr et al., 2016; Scheidelaar et al., 2016). At neutral to alkaline pH, deprotonation of the maleic acid groups promotes aqueous solubility, whereas the styrene moieties mediate hydrophobic interactions with lipid bilayers. Upon interaction with membranes, SMA can insert into the bilayer and induce its fragmentation into styrene–maleic acid lipid particles (SMALPs), in which a patch of lipid bilayer is stabilized by the amphipathic polymer (Dörr et al., 2016; Scheidelaar et al., 2016; Kuyler, 2025).

SMALPs are typically discoidal particles with diameters of approximately 10 nm, although their size and dispersity depend on polymer composition and experimental conditions. Factors affecting SMALP formation include the styrene-to-maleic-acid ratio, polymer conformation, pH, membrane phase state, and polymer-to-lipid ratio, with insufficient polymer favouring larger and more heterogeneous lipid assemblies (Dörr et al., 2016; Scheidelaar et al., 2016; Dominguez Pardo et al., 2017; Tang et al., 2026). SMA can also directly solubilize membrane proteins from biological membranes by excising patches of the surrounding lipid bilayer, thereby forming protein-containing SMALPs without the use of conventional detergents (Dörr et al., 2016; Lee et al., 2016). The dimensions of protein-containing SMALPs are influenced not only by the properties of the polymer and lipid bilayer, but also by the size and oligomeric state of the incorporated membrane protein.

It has been demonstrated that SMALPs support rapid lipid exchange (Cuevas, 2017), suggesting an opportunity to transfer membrane proteins between lipid environments and providing a physical basis for SMT. Lipids in SMA nanodiscs exchange through both aqueous monomer diffusion and, more rapidly, collision-mediated transfer. The latter has been attributed to the flexibility of the SMA belt, which permits transient formation of a hydrophobic continuum between interacting membrane patches, thereby lowering the kinetic barrier for lipid transfer without requiring complete exposure of the hydrocarbon chains to water (Cuevas, 2017). Thus, SMA facilitates exchange by making otherwise unfavorable kinetic pathways accessible, while the equilibrium distribution of lipids ultimately depends on their chemical potentials in the respective membrane environments. Although the mechanism of protein transfer is less well understood, addition of KcsA-containing SMA nanodiscs to planar lipid bilayers resulted in the appearance of functional KcsA channels, which was interpreted as spontaneous fusion of individual nanodiscs with the bilayer (Dörr et al., 2014). The molecular details of this transfer process, however, were not directly resolved.

Here, we exploit this property to develop the Synthetic Membrane Transfer (SMT), a new method for reconstituting transmembrane proteins into phospholipid GUVs under mild and physiologically relevant conditions. Beyond enabling functional reconstitution, a central motivation for developing SMT is to combine protein insertion with controlled membrane shaping. GUV-based approaches have been widely used to study curvature sensing by membrane-associated proteins, notably through tube-pulling assays (Morlot, 2012; Prévost, 2017; Schöneberg, 2018) and, more recently, through the Synthetic Membrane Shaper (SMS) method (De Franceschi, 2022; De Franceschi, 2024). In the SMS, GUVs produced by inverted emulsion are reshaped into stomatocyte or dumbbell geometries by DNA nanostructures, generating narrow membrane necks with a well-defined catenoid (saddle-shaped) geometry. Under hyperosmotic conditions, the reduced vesicle volume provides excess membrane area for deformation, while cholesterol-functionalized DNA nanostars generate membrane asymmetry and spontaneous curvature. The cholesterol-linked oligonucleotides act primarily through a wedge effect, while the membrane-bound DNA nanostars further promote curvature through molecular crowding, thereby stabilizing deformed geometries such as dumbbells and stomatocytes (De Franceschi, 2022).

While many membrane-associated proteins are known to sense or generate curvature (Prévost, 2015, Bertin, 2020; Jukic, 2022; De Franceschi, 2018), direct evidence for curvature sensing by transmembrane proteins remains scarce. A few transmembrane proteins, such as tetraspanins (Dharan, 2022) and the potassium channel KvAP (Aimon, 2014), have been reported to preferentially localize to curved membranes, but systematic reconstitution-based studies are rare, and evidence for sensing of catenoid-shaped membrane necks is absent. This gap is striking given the prevalence of transmembrane proteins implicated in catenoid-shaped, highly curved membrane neck regions, including nuclear pore components (Funakoshi, 2011; Amm, 2023), viral pore-forming proteins (Huang, 2024, den Boon, 2024), bacterial chemoreceptors (Strahl, 2015), DNA translocases (Chan, 2022), and subunits of the mitochondrial contact site and organizing system (MICOS). MICOS is an evolutionarily conserved, multi-subunit complex that has been shown to govern the formation and maintenance of mitochondrial crista junctions featuring a funnel or slit-like architecture (Daumke, 2025). It consists of at least six distinct subunits which are mostly small, membrane-embedded proteins of the inner mitochondrial membrane. Mic10 is thought to form a small hairpin-like structure in the inner mitochondrial membrane which oligomerizes in arc-like structures to stabilize the high curvature of the crista neck (Barbot, 2015; Bohnert, 2015). However, direct experimental tests of this hypothesis are lacking.

Here, we combine SMT and SMS in a single-step protocol to achieve reconstitution of transmembrane proteins in GUVs that are simultaneously sculpted into complex catenoid geometries (Figure 1A). This integrated platform enables direct interrogation of curvature-dependent sorting of transmembrane proteins in a minimal and well-controlled system. Using this approach, we show that the MICOS subunit Mic10 senses membrane curvature and becomes enriched at catenoid-shaped necks. Our results establish a general strategy for studying transmembrane proteins at regions of complex membrane curvature and open the door to reconstituting larger curvature-stabilizing protein assemblies in synthetic membranes.

**Figure 1.**
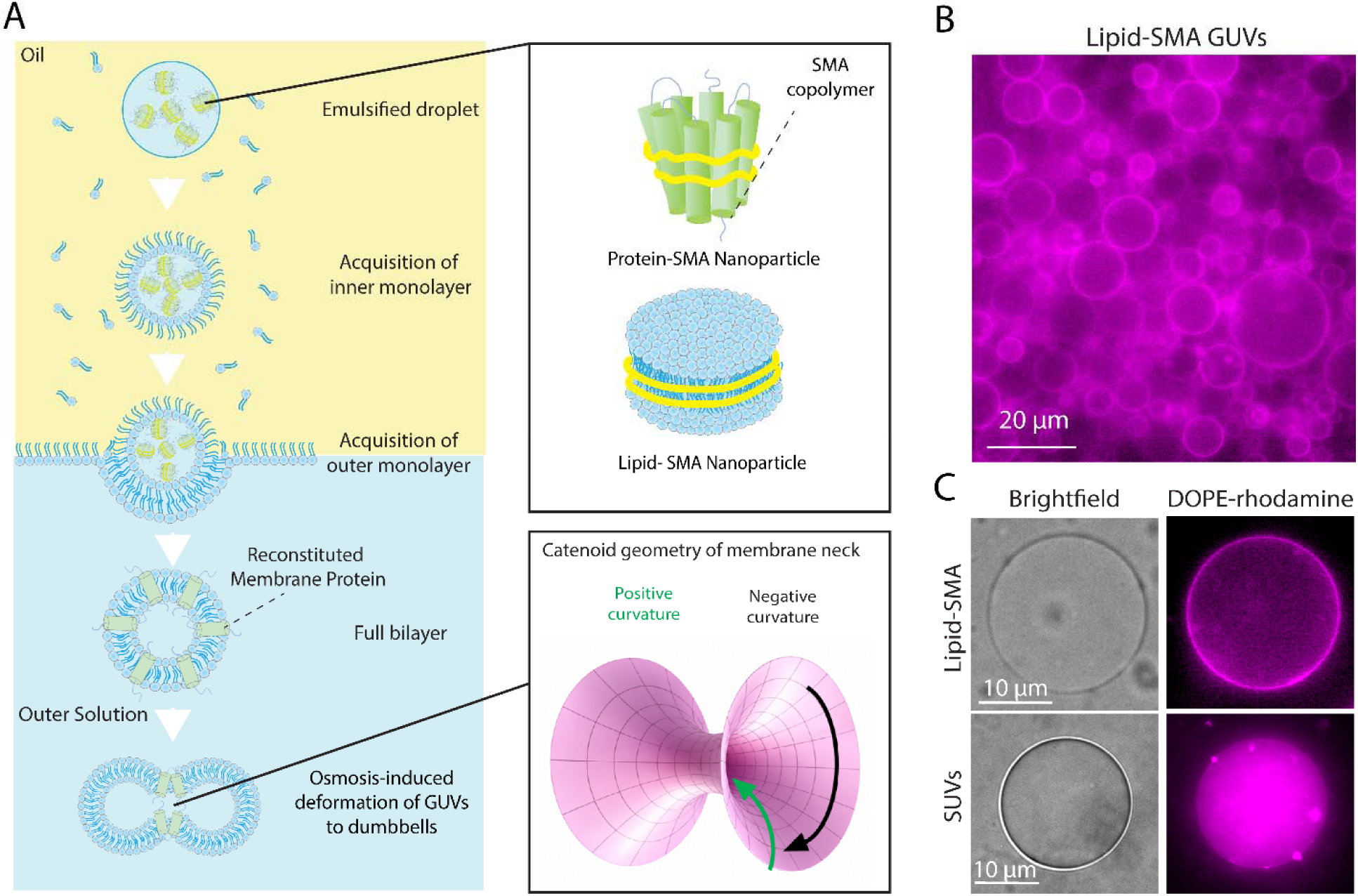
Lipid transfer from SMA nanoparticles to GUV membranes via Synthetic Membrane Transfer (SMT). (A) Schematic of the SMT approach. The inner aqueous solution containing lipid-SMA or protein-SMA is emulsified in a lipid-containing oil phase. The molecular organization of lipid-SMA and protein-SMA shown here is based on structures described in the available literature. As droplets traverse the oil and oil/outer buffer interface, they sequentially acquire inner and outer lipid monolayers to form a bilayer. During this process, lipid-SMA and protein-SMA interact with the nascent membrane and transfer to the GUV bilayer occur. The higher osmolarity of the outer solution and the presence of DNA nanostars induce deformation into dumbbell geometries with catenoid-shaped membrane necks. The directions of positive and negative curvature in the catenoid geometry are indicated. The intermediate steps involved in membrane deformation into dumbbell geometries are omitted for clarity. (B) Representative field of view of GUVs generated by the SMT. Membrane fluorescence (magenta) originates exclusively from Rhodamine-labeled lipids initially present in lipid-SMA. (C) Example of GUV generated by the SMT, in which fluorescent lipids have been transferred from SMA to the GUV membrane. In contrast, inclusion of fluorescent lipids in the form of SUVs in the inner buffer does not result in lipid transfer.

## Results

### Establishing SMT-Mediated Lipid Transfer into GUV Membranes

The Synthetic Membrane Transfer (SMT) approach exploits the intrinsic propensity of styrene–maleic acid (SMA) copolymers to exchange lipids and membrane-embedded material, thereby enabling transfer of phospholipids and integral membrane proteins (IMPs) into GUV membranes without detergents or fusogenic agents. Hereafter, we refer to SMA-embedded lipids and proteins as lipid-SMA and protein-SMA, respectively. As a first step toward IMPs reconstitution under curvature-controlled conditions, we tested whether lipid-SMA could be incorporated into GUV membranes during vesicle formation.

In SMT, GUVs are generated by inverted emulsion: aqueous droplets containing the inner buffer are emulsified in an oil phase containing lipids and subsequently transferred on top of an outer aqueous solution. As droplets sediment by gravity, they acquire first the inner and then the outer lipid monolayer upon crossing the oil/outer buffer interface, forming a bilayer. Simultaneously, cholesterol-functionalized DNA nanostars present in the outer solution bind to the nascent GUVs and, under hyperosmotic conditions, facilitate their reshaping into dumbbell geometries (De Franceschi, 2022) (Figure 1A). Lipid-SMA and protein-SMA are included in the inner buffer, such that lipid/IMPs transfer, GUV formation, and membrane reshaping occur on overlapping timescales: this is a key feature for reconstituting curvature-sensitive transmembrane proteins into defined membrane regions.

To assess lipid transfer, we prepared lipid-SMA particles containing fluorescent lipids. Inclusion of lipid-SMA in the inner buffer yielded GUVs with strong fluorescence colocalizing with the membrane (Figure 1B). In contrast, when Small Unilamellar Vesicles (SUVs) carrying the same fluorescent lipid were added at comparable concentrations, fluorescence was either confined to the lumen or appeared as aggregates (Figure 1C and Figure S1A), indicating inefficient lipid transfer.

These results are consistent with lipids incorporation into GUV membranes and define the physicochemical framework required for subsequent reconstitution of functional IMPs during membrane shaping.

### Functional Reconstitution of aquaporin-12 by SMT

Having established SMT-mediated lipid transfer into GUV membranes, we next asked whether this approach enables functional reconstitution of IMPs. As a proof-of-principle, we selected aquaporin-12 (AQP12), a well-characterized water channel (Bjørkskov, 2017). Purified GFP-tagged AQP12 was incorporated into SMA by detergent exchange using Bio-Beads. Upon SMT-mediated reconstitution, GUVs displayed a homogeneous GFP signal along the GUV rim (Figure 2A, B), consistent with AQP12 incorporation into the bilayer. To assess whether AQP-12 was functionally reconstituted, we exploited the ability of aquaporins to transport hydrogen peroxide. HLOL transport has been demonstrated for several other aquaporins, including AQP1 and AQP8, with permeability varying substantially among isoforms (Almasalmeh, 2014). However, HLOL transport by AQP12 has, to our knowledge, not previously been quantitatively characterized. GUVs were generated in the presence of a fluorescent HLOL sensor in both inner and outer solutions, and HLOL was introduced into the observation chamber by microfluidic perfusion.

**Figure 2.**
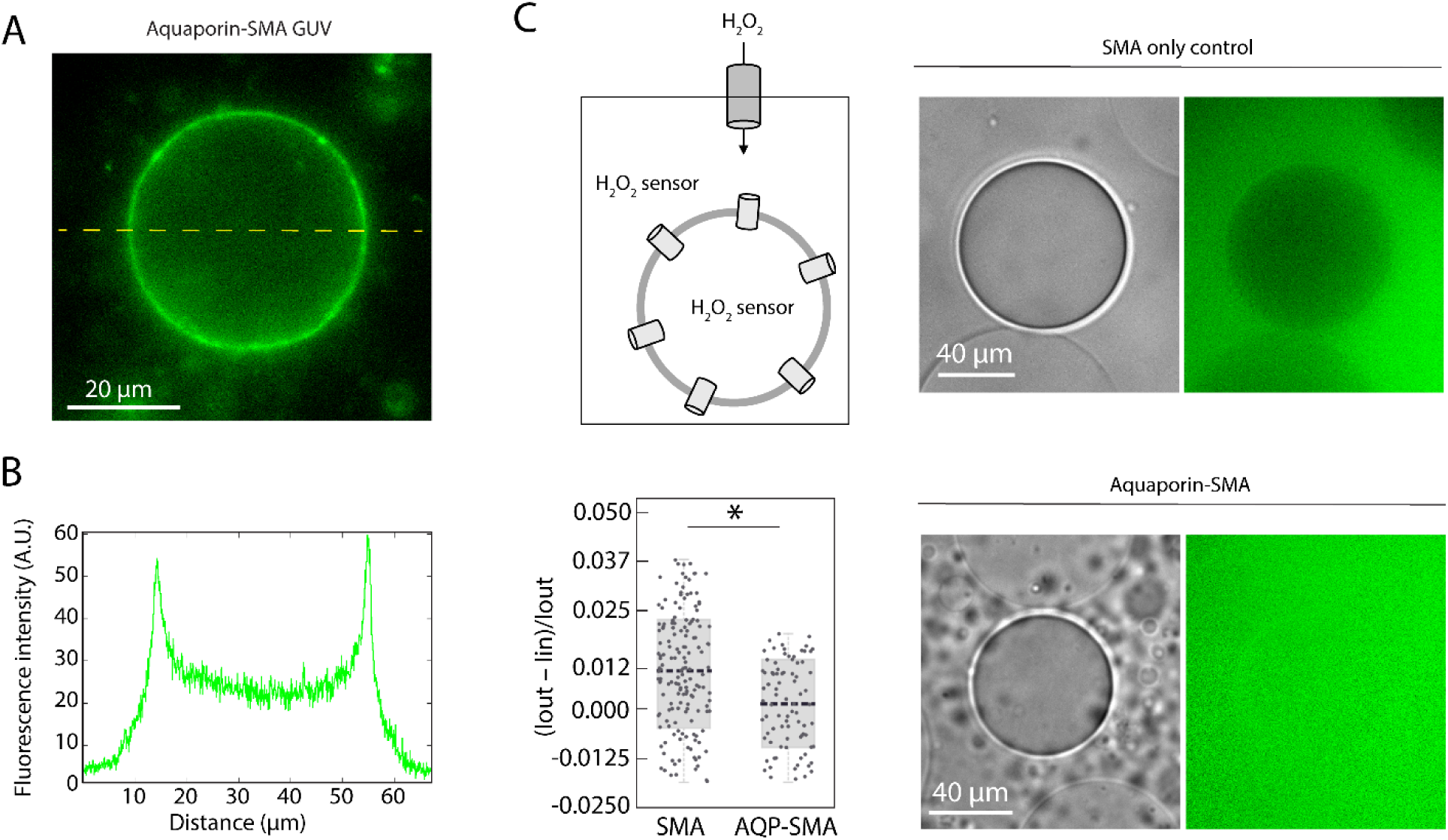
Functional reconstitution of AQP12 in GUVs by SMT. (A): Representative GUV with membrane-localized AQP12–SMA. The fluorescence from the GFP tag is displayed in green. (B): Line-scan analysis of the GUV shown in (A). (C) H□O□ influx assay. Representative images of GUVs containing AQP12–SMA or SMA-only control. Fluorescence from the sensor (in green) reports H□O□ permeation into the lumen. GUVs were incubated with H□O□ for 30 min before imaging. The quantification of H□O□ permeation, expressed as I_n,diff_ = (I_out_ – I_in_)/I_out_ (see Methods), shows increased H□O□ permeability compared to SMA-only controls. Values above zero indicate lower permeability. (Wilcoxon–Mann–Whitney test, two-sided, p = 0.0012; n = 108 for AQP12–SMA, n = 200 for control; 3 independent experiments).

Proteo-GUVs containing AQP12 exhibited a significantly higher fluorescent signal in the lumen than SMA-only controls (Figure 2C), demonstrating facilitated HLOL transport across the membrane. These data indicate that AQP12 is functionally reconstituted by SMT and, to our knowledge, provide the first direct demonstration by bottom-up reconstitution in GUVs that AQP12 transports HLOL. Crucially, this establishes SMT as a platform compatible with the subsequent characterization of geometry-dependent protein sorting during GUV shaping, which we address next.

### Curvature-Dependent Sorting of Mic10 at Catenoid Membrane Necks

To directly probe curvature sensing by transmembrane proteins, we focused on Mic10, a small membrane-embedded hairpin protein and a core component of the mitochondrial contact site and organizing system (MICOS) (Stephan, 2020). Mic10 is thought to oligomerize into arc-like assemblies that stabilize the extreme curvature of mitochondrial crista junctions (Barbot, 2015; Bohnert, 2015, Stephan, 2024), yet direct experimental evidence for curvature sensing by Mic10 in a minimal membrane system has been lacking.

Mic10 was recombinantly produced as a split-GFP fusion protein and reconstituted in SMA. Upon SMT-mediated transfer into GUV membranes under isotonic conditions, Mic10 formed discrete clusters that diffused laterally along the membrane (Figure 3A, B; Movie 1). We next combined Mic10 reconstitution with simultaneous membrane deformation by performing SMT under hyperosmotic conditions in the presence of cholesterol-functionalized DNA nanostars, which reshape GUVs into dumbbell geometries featuring catenoid necks (Figure 3C). Under these conditions, Mic10 clusters localized to the neck region connecting the two lobes of the dumbbell-shaped GUVs (Figure 3D, Figure S2; Movies 2 and 3). Protein enrichment at membrane necks can be quantified using the sorting ratio, a parameter that compared the fluorescence intensity of the protein at the neck normalized to that of protein bound on the relatively flat liposome membrane (De Franceschi, 2022). For Mic10, the sorting ratio was 4.19 ± 3.29 (Mean ± SD, n = 29), indicating robust and reproducible localization at neck. FRAP analysis confirmed that mic10 localizes at the neck of dumbbell liposomes connected by an open membrane neck (Figure S3). We interpret the observed neck enrichment as evidence consistent with functional membrane insertion of Mic10, since curvature-dependent localization is expected to require its appropriate incorporation into the membrane.

**Figure 3.**
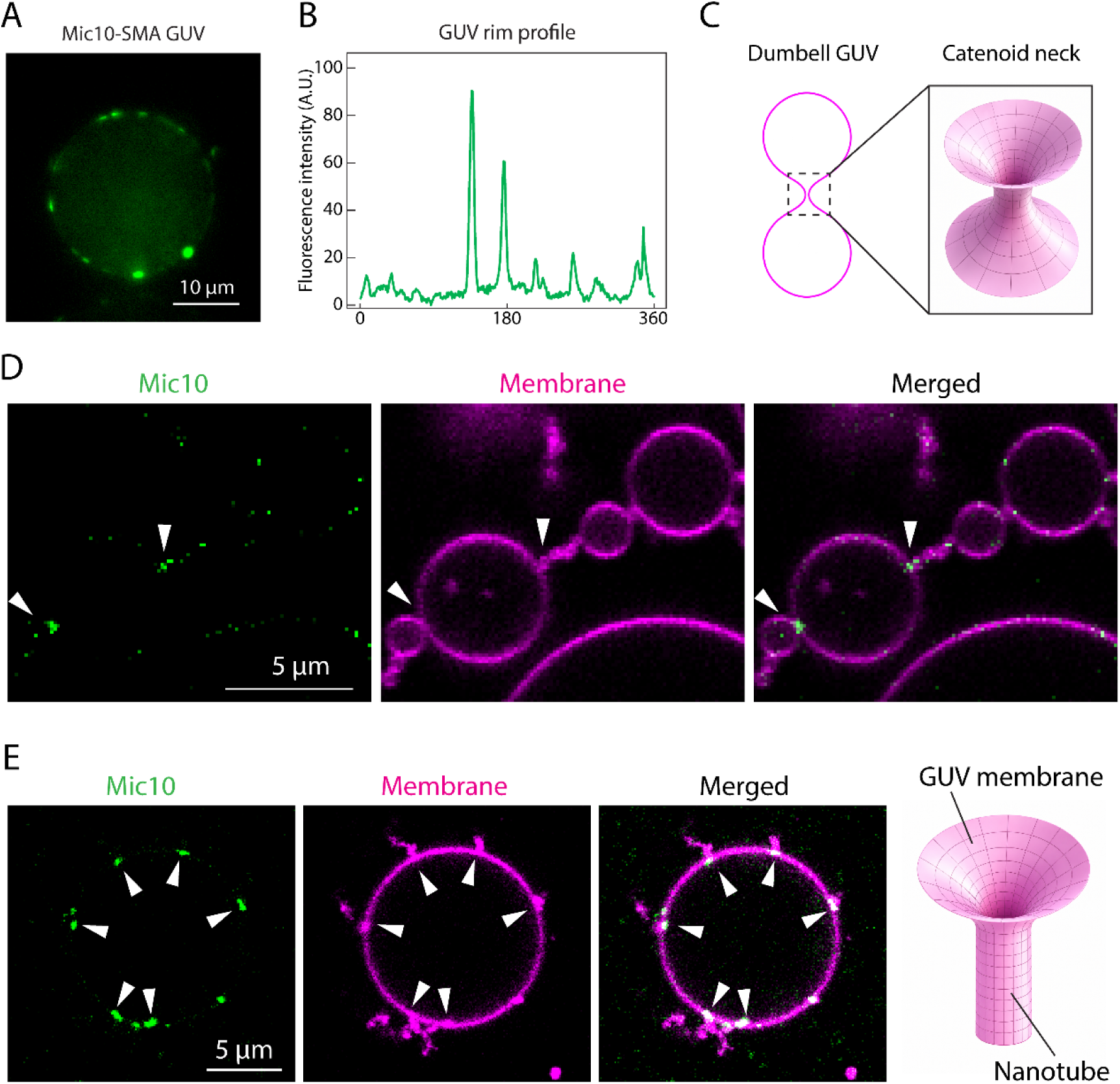
Curvature-dependent sorting of Mic10 at catenoid membrane necks. (A) Representative spherical GUV with membrane-localized clusters of Mic10–SMA. (B) Rim intensity profile of the GUV in (A), showing discrete peaks corresponding to Mic10 clusters. (C) Schematic illustrating the catenoid membrane geometry at the neck of dumbbells GUVs. (D) Confocal images of dumbbell-shaped GUVs formed under curvature-inducing conditions. Mic10 forms clusters that are strongly enriched at catenoid membrane necks (arrowheads). Membranes are labeled with DOPE–Rhodamine (magenta). (E) Confocal images of spherical GUVs displaying membrane nanotubes. Mic10 clusters localize at nanotube necks (arrowheads), consistent with curvature-dependent enrichment. Membranes are labeled with DOPE–Rhodamine (magenta). The schematic illustrates the geometry of the membrane forming the nanotubes.

Consistent with curvature-dependent localization, Mic10 also accumulated at the base of membrane nanotubes emanating from GUVs (Figure 3E; Movie 4), which arose spontaneously during the SMT procedure under the hyperosmotic conditions used to induce membrane deformation. These nanotubes match the membrane geometry of cristae junctions, where Mic10 was shown to localize (Kondadi, 2020). These observations closely parallel the behavior of well-established soluble curvature-sensing proteins such as ESCRT-III components (De Franceschi, 2018; Schöneberg, 2018; Souza, 2025), but here arise from a transmembrane protein.

Together, these results provide direct evidence that Mic10 senses and preferentially localizes to membranes of saddle-shaped curvature. More broadly, they establish the SMT as a powerful bottom-up platform to investigate the assembly of transmembrane protein complexes at regions of complex membrane geometry.

### Versatility of SMT for Co-Encapsulation and Membrane Protein Reconstitution

To assess the applicability of SMT for bottom-up synthetic biology, we next examined whether transmembrane protein reconstitution could be combined with co-encapsulation of macromolecules commonly used in synthetic cell models. A key requirement for such systems is the ability to control both lumenal composition and membrane functionality within the same compartment.

Using the SMT protocol, we reconstituted Mic10 upon co-encapsulation of representative macromolecules of different classes, including 10 kDa dextran and a 22-mer single-stranded DNA (ssDNA) in dumbbell-shaped GUVs. In these conditions, Mic10 retained its ability to form clusters and exhibited robust enrichment at catenoid membrane necks (Figure 4A, B), indicating that curvature sensing is preserved in the presence of lumenal macromolecular crowding.

**Figure 4.**
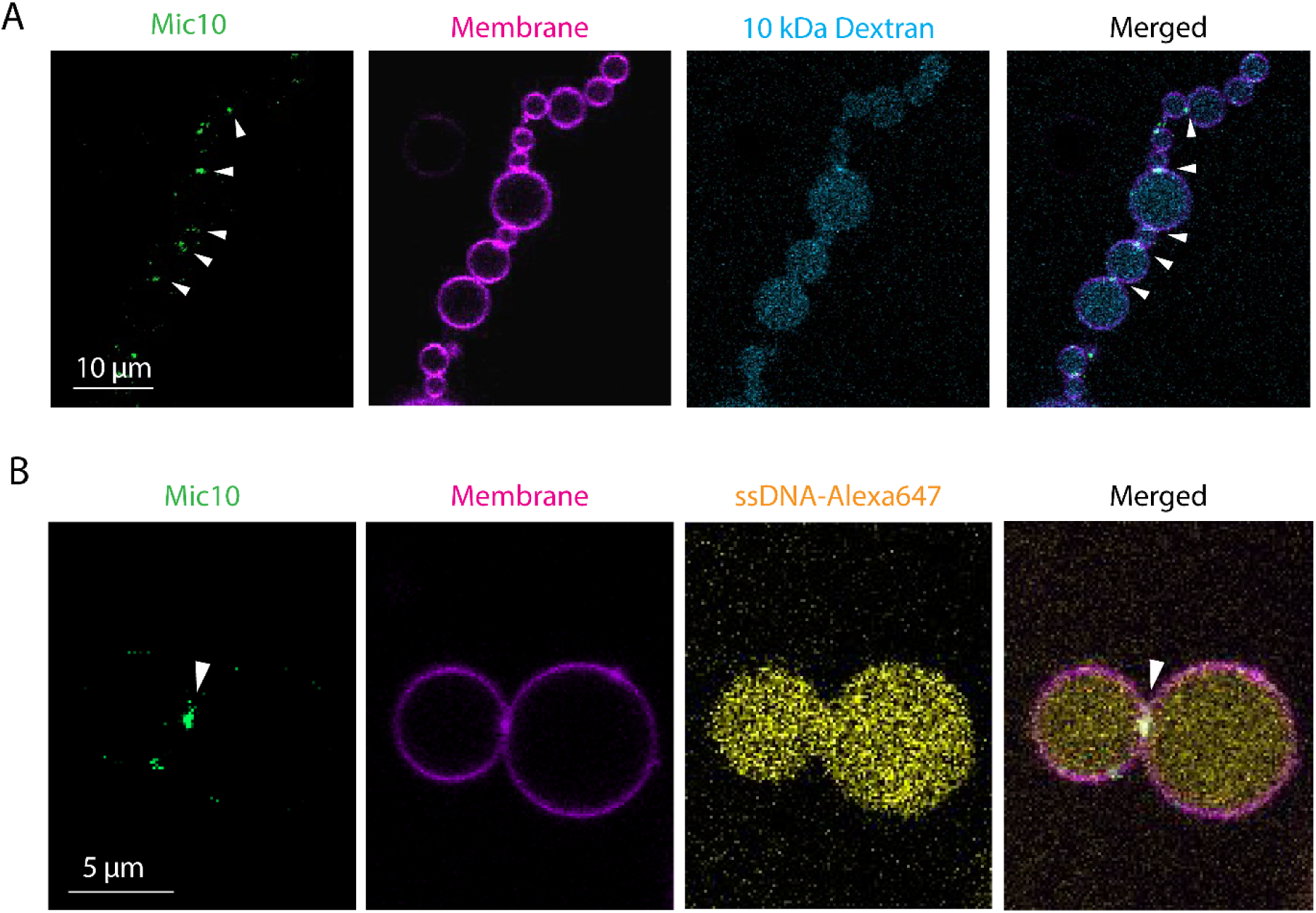
SMT enables simultaneous macromolecule encapsulation, membrane protein reconstitution, and curvature-dependent sorting. (A) Representative images of dumbbell-shaped GUVs containing lumenal 10 kDa fluorescent dextran with Mic10 reconstituted under curvature-inducing conditions. Mic10 retains clustering and enrichment at catenoid membrane necks (arrowheads) in the presence of encapsulated dextran. (B) Representative images of dumbbell-shaped GUVs containing a lumenal fluorescent ssDNA oligo with Mic10 reconstituted under curvature-inducing conditions. Mic10 retains clustering and enrichment at catenoid membrane necks (arrowhead) in the presence of encapsulated ssDNA.

Together, these results demonstrate that SMT enables simultaneous transmembrane protein reconstitution, membrane shaping, and macromolecular encapsulation in a single-step process. This capability establishes SMT as a versatile platform for bottom-up reconstitution of membrane-associated processes in geometrically defined compartments.

### Conclusions

We have developed Synthetic Membrane Transfer (SMT), a novel approach for detergent-free reconstitution of transmembrane proteins in GUVs that can be directly combined with controlled membrane shaping. The SMT enables protein insertion under mild conditions while remaining compatible with inverted emulsion protocols, thereby allowing simultaneous control of membrane composition, lumenal content, and membrane geometry.

Using the SMT, we achieved functional reconstitution of structurally diverse transmembrane proteins, including aquaporin-12 (AQP12) and the membrane-spanning hairpin protein Mic10. While aquaporins have been functionally characterized in nanoscale liposomes (Bjørkskov, 2017), functional reconstitution in GUVs has remained challenging. Here, we show by direct bottom-up reconstitution that AQP12 is capable of transporting HLOL in GUVs, extending functional studies of aquaporins to cell-sized compartments.

A central result of this work is the ability to probe curvature-dependent behavior of transmembrane proteins in a minimal system. In the absence of imposed curvature, Mic10 forms dynamic membrane-associated clusters, consistent with its propensity to oligomerize and remodel membranes. Upon induction of dumbbell geometries, Mic10 becomes strongly enriched at catenoid membrane necks, which recapitulate the saddle-shaped geometry of mitochondrial crista junctions. To our knowledge, this represents the first direct demonstration of a transmembrane protein preferentially localizing to a catenoid membrane geometry in a reconstituted system.

Notably, the membrane necks generated in this system have diameters in the range of 25–130 nm (De Franceschi, 2022), comparable to physiological curvature scales. This enables direct investigation of nanoscale curvature-dependent processes within micron-sized compartments, effectively bridging length scales that are typically difficult to access simultaneously.

Beyond curvature sensing, SMT is compatible with co-encapsulation of macromolecules and supports simultaneous membrane protein reconstitution, membrane shaping, and lumenal loading in a single step. This combination of features makes SMT well suited for bottom-up reconstitution of membrane-associated processes in geometrically defined environments.

This approach opens the door to reconstitution of multiprotein assemblies that operate at membrane necks, including the MICOS complex (Barbot, 2015; Bohnert, 2015), nuclear pore components (Funakoshi, 2011; Amm, 2023), viral RNA-transport machinery (Huang, 2024), and bacterial division machinery (Du, 2019) (Figure 5), which have remained challenging to access using existing methods.

**Figure 5.**
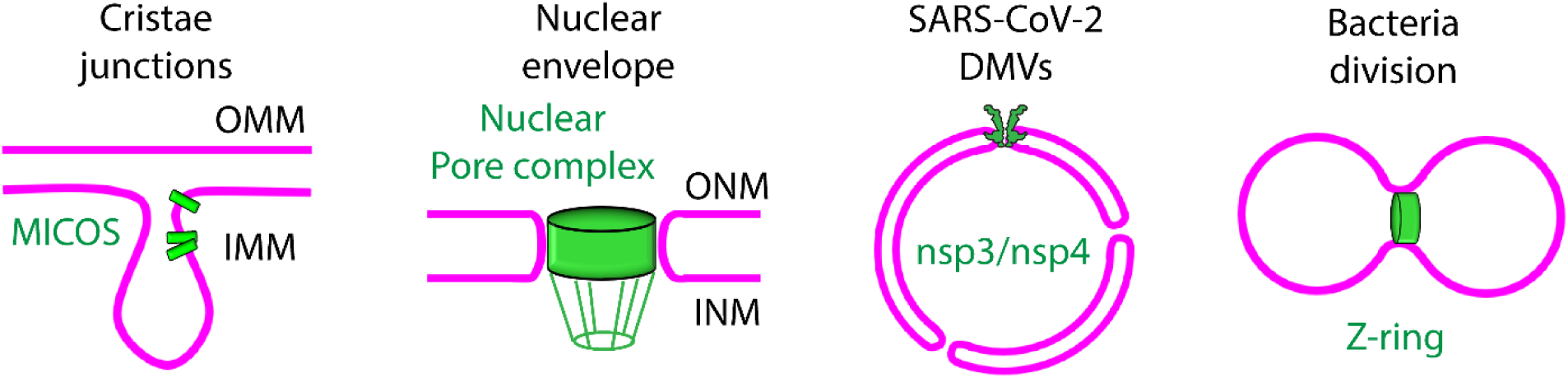
Schematic illustrating representative biological systems in which transmembrane proteins assemble at catenoid membrane necks. These include mitochondrial crista junctions (MICOS), nuclear pore complexes, SARS-CoV-2 double-membrane vesicles hosting nsp3/nsp4 pores, and bacterial division sites (Z-ring). Membranes are shown in magenta and protein assemblies in green. OMM, outer mitochondrial membrane; IMM, inner mitochondrial membrane; ONM, outer nuclear membrane; INM, inner nuclear membrane; DMV, double-membrane vesicle.

## Methods

### Reagents

Glucose (G7021), MgCl_2_ (M8266), silicone oil (317667), mineral oil (M3516-1L), and Optiprep (60% (w/v) iodixanol in water (D1556) were purchased from Sigma-Aldrich. Tris-HCl (10812846001) was purchased from Roche. DOPC (1,2-dioleoyl-sn-glycero-3-phosphocholine) (850375), EggPC (840051C), DOPE-Rhodamine (1,2-dioleoyl-sn-glycero-3-phosphoethanolamine-N-(lissamine rhodamine B sulfonyl) (ammonium salt) (810150C), and DOPS (1,2-dioleoyl-sn-glycero-3-phospho-L-serine (sodium salt) (840035C) were purchased from Avanti Lipids. Lipids were stored and resuspended in anhydrous chloroform (288306, Sigma-Aldrich). Bovine serum albumin was purchased from SERVA (Lot 240027). 10 kDa dextran-647 was purchased from Invitrogen (D22914). SMA (SMALP 200) was purchased from Aurorium. n-dodecyl-β-D-maltopyranoside (D310S) was purchased from Anatrace. The OxiVision™ Green hydrogen peroxide sensor (AATB21505) was purchased from AAT Bioquest.

### Solutions

In this manuscript, we refer to the “inner solution” as the solution present inside the lumen of the GUVs, that is used to generate droplets. This solution was composed of 50 mM Tris pH 7.5, 30% Optiprep. The “outer solution” is the solution present outside the GUVs: this was composed of 50mM Tris pH 7.5, to which glucose was added until reaching the desired osmolarity. The osmolarity of the solutions was measured using an Osmomat 3000 (Gonotec).

### Lipid-in-Oil Suspension and Droplet Preparation

Lipid-in-oil suspensions were prepared based on published protocols (Van de Cauter, 2021; De Franceschi, 2022) with some modifications. The lipid mix was composed of 90 mol% EggPC and 10 mol% DOPS. 0.05 mol% DOPE-Rhodamine was included when imaging GUVs with reconstituted Mic10. Lipids were mixed in chloroform. Under argon stream, the freshly prepared mixture of silicone and mineral oil was added to the lipids dropwise while vortexing at 1,400 rpm. The lipid-in-oil solution was finally vortexed at 2,800 rpm for 2 min and further sonicated in an ice bath for 15 min. Droplets were generated by mixing 10 µl of inner solution into 400 µl of lipid-in-oil suspension. The suspension was emulsified by vigorous pipetting for 30 seconds and subsequently added on top of the 200 µl of outer solution in the observation chamber. The sample was incubated for 1 hour at 23 °C to allow GUVs to sink and subsequently imaged.

### Protein cloning, Expression, Purification, and Labeling

#### Aquaporin-12

The plasmid encoding for aquaporin-12 fused to an N-terminal EGFP was previously described (Bjørkskov et al., 2017). Yeast cells from 4 L cultures were harvested at 2,000 g for 10 minutes at 4 °C. Cells were lysed by glass bead homogenization in ice cold Lysis buffer (25 mM Tris pH 8.0, 1 mM EDTA, 1 mM EGTA, 10% glycerol, 250 mM NaCl) containing 1 mM PMSF and 1 μg/ml of leupeptin, pepstatin and chymostatin as described previously. The cell lysate was centrifuged at 4 °C at 3,000 g for 30 minutes and the supernatant was exposed to ultracentrifugation at 120,000 g for 2 hours. Crude membrane pellets were resuspended in lysis buffer containing protease inhibitors and kept at −80 °C until use. Crude membranes were solubilized ON in a glass beaker at 4 °C by slow rotation on a magnetic stirrer at a protein concentration of 5 mg/ml and 1% DDM (Glycon, Germany) + 0.33% cholesterol-hemisuccinate (CHS) (Anatrace, USA). Solubilized protein was isolated by ultracentrifugation at 4 °C for 1 hour at 120,000 g and diluted with two volumes of lysis buffer containing protease inhibitors but no detergent. The diluted solubilized proteins were incubated overnight with Indigo resin (Cube Biotech, Germany) in a glass beaker with slow magnetic stirring. Next morning, the beaker content was transferred to an Econo disposable column (Biorad, USA). After collecting the run-through, the column was washed with Lysis buffer containing 10 mM, 30 mM, 100 mM, 250 mM or 500 mM imidazole. All buffers contained detergent at a concentration of 0.4% DDM and 0.0136 % CHS, 1 mM PMSF and 1 μg/ml of leupeptin, pepstatin and chymostatin. Fluorescence in each fraction was quantified using a spectrofluorometer (Fluoroskan Ascent, Thermo Fisher Scientific) using buffer as a blank. Excitation was at 485 nm and emission at 520 nm. The 250 mM eluate was concentrated on a Vivaspin filter with a 10 kDa cutoff.

#### Mic10

To obtain the protein in a functional form, we pursued an approach described for stabilizing the flexible termini of a vitamin K epoxide reductase (Liu, 2021). Thus, we recombinantly produced and purified Drosophila Mic10b (Stephan, 2024) in DDM as a split GFP-fusion protein, in which both the N- and C-termini were fused to the two moieties of split-GFP. The two fragments of split-superfolderGFP were fused to the N- and C-terminus of *Drosophila melanogaster* MIC10b, respectively. The coding sequence of the fusion protein was synthesized and incorporated into pET11a via NdeI/BamHI by GenScript:

Atgtcaaaaggagaagagctatttacaggggtagtgccgattctggtggagctggatggagacgtcaacggtcataagttcagcgtgagaggcgaggg cgaaggtgatgcaaccaatggcaagttgaccttaaagttcatctgcaccaccggtaaactgccggttccgtggccaacattggttacgactctgacctatg gtgttcaatgcttttctcgctacccggaccacatgaaacgccatgatttttttaagagcgctatgccggaaggctatgttcaagaacgtactatttcgttcaag gacgacggcacctacaaaacccgtgcggaagttaaattcgagggcgacaccttggtgaaccgtatcgagctcaagggtatcgatttcaaagaagatggt aatattcttggtcacaaactggagtacaacatgaccagccgtgataacatttttgaggaaaaaatttgtaatcgtctggaccactgcgttagtgatgtgctgat taaaggttgtggtggtgtcatcatcggtagcgccgttagcttcctgatcctgaagcgccgtgcgtggccagtttggctgggtgcgggttttggcatgggtat cgcctatcgtacgtgcgaaaaggacctgaactctctgaagaacagccataacgtgtatatcacggcggataaacagaaaaacggcatcaaagctaatttt aagattcgtcacaacgtcgaggatggctccgtgcaactggcggatcactaccagcagaataccccgattggcgacggaccggtactccttccggacaa ccattacctgtccactcagagcgttttgtccaaagacccgaatgaaaagcgcgaccacatggtgctgctggagttcgtgaccgcggcaggcatcacccat caccatcaccaccaccactaa.

The expression plasmid was freshly transformed into *E. coli* C41 (DE3) cells and protein expression was carried out in terrific broth containing 100 µg/ml ampicillin. The cultures were grown at 37 °C and 80 rpm until OD_600_ reached 0.8, protein expression was subsequently induced by the addition of 300 µM isopropyl β-D-1-thiogalactopyranoside (IPTG), and cells were incubated at 20 °C for another 16 hours. The next day, the cells were sedimented at 4,000 g, collected, and frozen at -20 °C.

Cells expressing split-GFP-Mic10 were diluted in 50 mM HEPES/NaOH pH 7.5, 500 mM NaCl and 1 mg/ml DNase I (Roche) before lysis using a microfluidizer (Microfluidics). To remove insoluble parts, the solution was centrifuged at 15,000 g, 4 °C for 20 min. A second centrifugation step at 130,000 g, 4 °C for 2 h was applied to pellet the membrane fraction of the *E. coli* cells and the resulting pellet was homogenized in resuspension buffer (50 mM HEPES/NaOH pH 7.5, 500 mM NaCl, 10 mM imidazole and 10% glycerol). To extract the membrane proteins, the homogenous solution was diluted in solubilization buffer (50 mM HEPES/NaOH pH 7.5, 300 mM NaCl, 10 mM imidazole, 2 mM DTT and 1% DDM) and incubated overnight at 4 °C under gentle stirring. The solution was centrifuged a second time at 130,000 g, 4 °C for 30 min. The resulting supernatant was further loaded onto a prepacked Ni^2+^ - Sepharose High Performance IMAC resin (Cytiva) containing gravity flow column charged with 100 mM nickel sulfate and equilibrated in Ni-IMAC wash buffer (50 mM HEPES/NaOH pH 7.5, 300 mM NaCl, 10 mM imidazole, 2 mM DTT and 0.2% DDM). The column was extensively washed using Ni-IMAC wash buffer and bound proteins were eluted with Ni-IMAC elution buffer containing 50 and 500 mM imidazole, respectively. Fractions containing split-GFP-Mic10 fusion protein were combined, concentrated, and subjected to size-exclusion chromatography (SEC) using a Superose 6 10/300 column (Cytiva) equilibrated in buffer SEC (20 mM HEPES/NaOH pH 7.5, 150 mM NaCl, 2 mM DTT and 0.03% DDM). Fractions containing pure protein were combined, concentrated to 13 mg/ml, flash frozen, and stored at -70°C.

### Lipid and Protein reconstitution in SMA

Lipid-SMA particles were generated as follows.

1 mg of lipids (89.999 mol% of eggPC + 10 mol% DOPS + 0.001 mol% of Rhodamine-DOPE) in choloroform were deposited in a glass vial and the chlorofom was removed by evaporation. The lipids were resuspended in 200 ul of buffer (150 mM NaCl + 50 mM Tris pH7.5 + 1 mM EDTA + 2 mM MgCl_2_). 15uL SMA (10%) were added and incubated for 30 minutes at 37C. The initially turbid lipid suspension clarified.

Protein-SMA particles were generated as follow.

AQP12: 6.6 ul of 10% SMA were added to 30 ul protein stock (at 2 mg/ml) in 0.1% DDM and incubated for 3 hours at 4 °C. Biobeads were added and the samples were incubated overnight shaking at 300 rpm and 4 °C to remove the DDM. After overnight incubation, the biobeads were removed and the sample was used for reconstitution in GUVs without further purification steps.

Mic10: 22.5 ul of 10% SMA were added to 10 ul protein stock at (13.8 mg/ml) in 0.1% DDM and incubated for 4 hours at 4 °C. Biobeads were added and the samples were incubated overnight shaking at 300 rpm and 4 °C to remove the DDM. After overnight incubation, the biobeads were removed and the sample was used for reconstitution in GUVs without further purification steps.

The concentrations reported for lipid-SMA and protein-SMA in the manuscript figures were calculated from their concentrations before detergent removal and corrected for the volume of SMA solution added. Bio-Beads were added as a semi-dry slurry and did not appreciably alter the sample volume. The final concentration of protein-SMA used in the inner buffer to generating GUVs were 113 nM for AQP12-SMA and 150 nM for Mic10-SMA.

### FRAP assay on GUVs

FRAP assay was performed using a confocal laser microscope (Zeiss LSM 910, 63× objective). To measure the recovery signal, frames were collected every at 1fps, starting right after the photobleaching event. Fluorescence images were analyzed using ImageJ.

### Quantification of fluorescent molecules influx in GUVs

Quantification was performed as described in (Fragasso, 2021). Regions of interest (ROIs) were selected both inside and immediately outside GUVs. The average fluorescent signal from the H_2_O_2_ sensor was measured in these ROIs and for each GUVs we calculated the normalized fluorescence intensity difference according to the formula: I_n,diff_ = (I_out_ – I_in_)/I_out_.

### Rim scan analysis

To analyze pixel intensity along the vesicle membrane, we developed a custom Python-based application using OpenCV and Matplotlib. The tool enables manual selection and interactive adjustment of elliptical or circular regions corresponding to vesicle membranes in grayscale microscopy images. Intensity profiles are calculated by sampling pixel values at 360 angular positions along the ellipse. For each angle, intensities are averaged over the specified rim thickness, e.g., 6 pixels used in this study, to reduce noise. Once the rim of a vesicle membrane is selected, the application plots intensity versus angular position and exports the data in CSV format for further analysis. This tool facilitates reproducible quantification of membrane-associated fluorescence or other signal distributed along vesicle contours and is available upon request.

### Quantification of sorting ratio

In dumbbell-shaped vesicles, protein signal intensity at neck region and adjacent membrane was measured using multiple line scans and average of maximum signal intensity was used for further calculation. The background signal intensity was also measured. The sorting ratio was calculated with the following formula: Sorting Ratio = (I_max(neck)_ – I_background_)/(I_max(membrane)_ – I_background_).

## Author Contributions

N.D.F conceptualized the research. H.M. and N.D.F. set up the SMT protocol. H.M. and N.D.F. performed the SMT and carried out the microscopy experiments. R.Z. wrote and applied the script for rim analysis. S.D.O.K. and P.A.P. purified aquaporin. T.B.B., T.S., S.J. and O.D. cloned and purified Mic10.

N.D.F. wrote the manuscript with input from all other authors.

## Acknowledgements

We thank Paul Beales for useful discussion. This research is part of the project No. 2022/45/P/NZ1/01565 co-funded by the National Science Centre and the European Union Framework Programme for Research and Innovation Horizon 2020 under the Marie Skłodowska-Curie grant agreement No. 945339. For the purpose of Open Access, the author has applied a CC-BY public copyright licence to any Author Accepted Manuscript (AAM) version arising from this submission. We acknowledge funding support from the NCN Sonata Bis 2022/46/E/NZ1/00160. T.S. acknowledges financial support from SFB1507 from Deutsche Forschungsgemeinschaft and Goethe University Frankfurt (SCALE - SubCellular Architecture of LifE). SJ acknowledges support from the European Research Council (ERCAdG No. 835102).

## Declaration of generative AI and AI-assisted technologies in the manuscript preparation process

During the preparation of this work, the authors used ChatGPT solely to improve the grammar, spelling, and readability of the manuscript. All scientific content, interpretations, and conclusions were generated and verified by the authors. The authors reviewed and edited the output as needed and take full responsibility for the content of the published article.

## Funding statement

This research is part of the project No. 2022/45/P/NZ1/01565 co-funded by the National Science Centre and the European Union Framework Programme for Research and Innovation Horizon 2020 under the Marie Skłodowska-Curie grant agreement No. 945339. For the purpose of Open Access, the author has applied a CC-BY public copyright licence to any Author Accepted Manuscript (AAM) version arising from this submission. We acknowledge funding support from the NCN Sonata Bis 2022/46/E/NZ1/00160, from SFB1507 from Deutsche Forschungsgemeinschaft and Goethe University Frankfurt (SCALE - SubCellular Architecture of LifE) and from the European Research Council (ERCAdG No. 835102).

## Conflict of interest disclosure

The authors declare that they have no known competing financial interests or personal relationships that could have appeared to influence the work reported in this paper.

## Notes

### Competing Interest Statement

The authors have declared no competing interest.

### Summary of Updates

This version of the manuscript has been revised to sharpen its focus on the reconstitution of integral membrane proteins in synthetic membranes with complex geometries, particularly their localization at catenoid membrane necks. The experimental results and discussion have been streamlined to emphasize the central methodological advance and its application to curvature dependent protein sorting. The text and figures have been reorganized accordingly to provide a more coherent and focused presentation of the principal findings.

## References

Aimon S, Manzi J, Schmidt D, Poveda Larrosa JA, Bassereau P, Toombes GE. Functional reconstitution of a voltage-gated potassium channel in giant unilamellar vesicles. PLoS One. 2011;6(10):e25529.

Aimon S, Callan-Jones A, Berthaud A, Pinot M, Toombes GE, Bassereau P. Membrane shape modulates transmembrane protein distribution. Dev Cell. 2014 Jan 27;28(2):212–8.

Alam S, Doherty E, Ortega-Prieto P, Arizanova J, Fets L. Membrane transporters in cell physiology, cancer metabolism and drug response. Dis Model Mech. 2023 Nov 1;16(11):dmm050404.

Almasalmeh A, Krenc D, Wu B, Beitz E. Structural determinants of the hydrogen peroxide permeability of aquaporins. FEBS J. 2014 Feb;281(3):647–56. doi: 10.1111/febs.12653.

Almendro-Vedia VG, Natale P, Mell M, Bonneau S, Monroy F, Joubert F, López-Montero I. Nonequilibrium fluctuations of lipid membranes by the rotating motor protein F1F0-ATP synthase. Proc Natl Acad Sci U S A. 2017 Oct 24;114(43):11291–11296. doi: 10.1073/pnas.1701207114.

Altamura E, Milano F, Tangorra RR, Trotta M, Omar OH, Stano P, Mavelli F. Highly oriented photosynthetic reaction centers generate a proton gradient in synthetic protocells. Proc Natl Acad Sci U S A. 2017 Apr 11;114(15):3837–3842. doi: 10.1073/pnas.1617593114.

Amm I, Weberruss M, Hellwig A, Schwarz J, Tatarek-Nossol M, Lüchtenborg C, Kallas M, Brügger B, Hurt E, Antonin W. Distinct domains in Ndc1 mediate its interaction with the Nup84 complex and the nuclear membrane. J Cell Biol. 2023 Jun 5;222(6):e202210059.

Barbot M, Jans DC, Schulz C, Denkert N, Kroppen B, Hoppert M, Jakobs S, Meinecke M. Mic10 oligomerizes to bend mitochondrial inner membranes at cristae junctions. Cell Metab. 2015 May 5;21(5):756–63.

Bertin A, de Franceschi N, de la Mora E, Maity S, Alqabandi M, Miguet N, di Cicco A, Roos WH, Mangenot S, Weissenhorn W, Bassereau P. Human ESCRT-III polymers assemble on positively curved membranes and induce helical membrane tube formation. Nat Commun. 2020 May 29;11(1):2663.

Biner O, Schick T, Müller Y, von Ballmoos C. Delivery of membrane proteins into small and giant unilamellar vesicles by charge-mediated fusion. FEBS Lett. 2016 Jul;590(14):2051–62.

Bohnert M, Zerbes RM, Davies KM, Mühleip AW, Rampelt H, Horvath SE, Boenke T, Kram A, Perschil I, Veenhuis M, Kühlbrandt W, van der Klei IJ, Pfanner N, van der Laan M. Central role of Mic10 in the mitochondrial contact site and cristae organizing system. Cell Metab. 2015 May 5;21(5):747–55.

Bjørkskov FB, Krabbe SL, Nurup CN, Missel JW, Spulber M, Bomholt J, Molbaek K, Helix-Nielsen C, Gotfryd K, Gourdon P, Pedersen PA. Purification and functional comparison of nine human Aquaporins produced in Saccharomyces cerevisiae for the purpose of biophysical characterization. Sci Rep. 2017 Dec 4;7(1):16899.

Chan H, Mohamed AMT, Grainge I, Rodrigues CDA. FtsK and SpoIIIE, coordinators of chromosome segregation and envelope remodeling in bacteria. Trends Microbiol. 2022 May;30(5):480–494. doi: 10.1016/j.tim.2021.10.002.

Cooper A, Vijayananda V, Pazzi J, Subramaniam AB. Combining Nanoscale Curvature and Polymer Osmotic Pressure for Efficient Giant Vesicle Assembly Under Physiological Conditions. Small Methods. 2025 Dec;9(12):e00664. doi: 10.1002/smtd.202500664.

Cuevas Arenas R, Danielczak B, Martel A, Porcar L, Breyton C, Ebel C, Keller S. Fast Collisional Lipid Transfer Among Polymer-Bounded Nanodiscs. Sci Rep. 2017 Apr 5;7:45875.

Daumke O, van der Laan M. Molecular machineries shaping the mitochondrial inner membrane. Nat Rev Mol Cell Biol. 2025 May 14.

den Boon JA, Nishikiori M, Zhan H, Ahlquist P. Positive-strand RNA virus genome replication organelles: structure, assembly, control. Trends Genet. 2024 Aug;40(8):681–693. doi: 10.1016/j.tig.2024.04.003.

De Franceschi N, Alqabandi M, Miguet N, Caillat C, Mangenot S, Weissenhorn W, Bassereau P. The ESCRT protein CHMP2B acts as a diffusion barrier on reconstituted membrane necks. J Cell Sci. 2018 Aug 3;132(4):jcs217968.

De Franceschi N, Pezeshkian W, Fragasso A, Bruininks BMH, Tsai S, Marrink SJ, Dekker C. Synthetic Membrane Shaper for Controlled Liposome Deformation. ACS Nano. 2022 Nov 28;17(2):966–78.

De Franceschi N, Barth R, Meindlhumer S, Fragasso A, Dekker C. Dynamin A as a one-component division machinery for synthetic cells. Nat Nanotechnol. 2024 Jan;19(1):70–76.

Dharan R, Goren S, Cheppali SK, Shendrik P, Brand G, Vaknin A, Yu L, Kozlov MM, Sorkin R. Transmembrane proteins tetraspanin 4 and CD9 sense membrane curvature. Proc Natl Acad Sci USA. 2022 Oct 25;119(43):e2208993119.

Dominguez Pardo, J. J., et al. (2017). Solubilization of lipids and lipid phases by the styrene–maleic acid copolymer. European Biophysics Journal 46, 91–101. DOI: 10.1007/s00249-016-1181-7.

Dörr, J. M. et al. (2016). The styrene–maleic acid copolymer: a versatile tool in membrane research. European Biophysics Journal 45, 3–21. DOI: 10.1007/s00249-015-1093-y.

Dörr JM, Koorengevel MC, Schäfer M, Prokofyev AV, Scheidelaar S, van der Cruijsen EA, Dafforn TR, Baldus M, Killian JA. Detergent-free isolation, characterization, and functional reconstitution of a tetrameric K+ channel: the power of native nanodiscs. Proc Natl Acad Sci U S A. 2014 Dec 30;111(52):18607–12. doi: 10.1073/pnas.1416205112.

Du S, Lutkenhaus J. At the Heart of Bacterial Cytokinesis: The Z Ring. Trends Microbiol. 2019 Sep;27(9):781–791.

Fragasso A, De Franceschi N, Stömmer P, van der Sluis EO, Dietz H, Dekker C. Reconstitution of Ultrawide DNA Origami Pores in Liposomes for Transmembrane Transport of Macromolecules. ACS Nano. 2021 Aug 24;15(8):12768–12779.

Funakoshi T, Clever M, Watanabe A, Imamoto N. Localization of Pom121 to the inner nuclear membrane is required for an early step of interphase nuclear pore complex assembly. Mol Biol Cell. 2011 Apr;22(7):1058–69.

Gaul V, Lopez SG, Lentz BR, Moran N, Forster RJ, Keyes TE. The lateral diffusion and fibrinogen induced clustering of platelet integrin αIIbβ3 reconstituted into physiologically mimetic GUVs. Integr Biol (Camb). 2015 Apr;7(4):402–11. doi: 10.1039/c5ib00003c.

Huang Y, Wang T, Zhong L, Zhang W, Zhang Y, Yu X, Yuan S, Ni T. Molecular architecture of coronavirus double-membrane vesicle pore complex. Nature. 2024 Sep;633(8028):224–231.

Kumar H, Mandal S, Yadav R, Gupta S, Meena H, Kadu M, et al. Bottom-up approach to explore alpha-amylase assisted membrane remodelling. Chemistry and Physics of Lipids. 2024;259:105374. doi: 10.1016/j.chemphyslip.2023.105374

Kondadi AK, Anand R, Hänsch S, Urbach J, Zobel T, Wolf DM, Segawa M, Liesa M, Shirihai OS, Weidtkamp-Peters S, Reichert AS. Cristae undergo continuous cycles of membrane remodelling in a MICOS-dependent manner. EMBO Rep. 2020 Mar 4;21(3):e49776. doi: 10.15252/embr.201949776.

Kuyler GC, Barnard E, Cunningham RD, Sibariboyi S, White L, Wessels I, Smith MP, Motloung B, Klumperman B. Amphiphilic Copolymers and Their Role in the Study of Membrane Proteins. J Phys Chem Lett. 2025 Jun 12;16(23):5784–5799.

Jukic N, Perrino AP, Humbert F, Roux A, Scheuring S. Snf7 spirals sense and alter membrane curvature. Nat Commun. 2022 Apr 21;13(1):2174.

Lee, S. C. et al. (2016). A method for detergent-free isolation of membrane proteins in their local lipid environment. Nature Protocols 11, 1149–1162. DOI: 10.1038/nprot.2016.070.

Li AZT, LiWang A, Subramaniam AB. Reconstitution of circadian clock in synthetic cells reveals principles of timekeeping. Nat Commun. 2025 Jul 21;16(1):6686. doi: 10.1038/s41467-025-61844-5.

Litschel T, Schwille P. Protein Reconstitution Inside Giant Unilamellar Vesicles. Annu Rev Biophys. 2021 May 6;50:525–548.

Liu S, Li S, Shen G, Sukumar N, Krezel AM, Li W. Structural basis of antagonizing the vitamin K catalytic cycle for anticoagulation. Science. 2021 Jan 1;371(6524):eabc5667.

Morlot S, Galli V, Klein M, Chiaruttini N, Manzi J, Humbert F, Dinis L, Lenz M, Cappello G, Roux A. Membrane shape at the edge of the dynamin helix sets location and duration of the fission reaction. Cell. 2012 Oct 26;151(3):619–29.

Nakamura S, Minamino T. Structure and Dynamics of the Bacterial Flagellar Motor Complex. Biomolecules. 2024 Nov 22;14(12):1488.

Olivi L, Berger M, Creyghton RNP, De Franceschi N, Dekker C, Mulder BM, Claassens NJ, Ten Wolde PR, van der Oost J. Towards a synthetic cell cycle. Nat Commun. 2021 Jul 26;12(1):4531.

Prévost C, Tsai FC, Bassereau P, Simunovic M. Pulling Membrane Nanotubes from Giant Unilamellar Vesicles. J Vis Exp. 2017 Dec 7;(130):56086.

Ripoll L, von Zastrow M, Blythe EE. Intersection of GPCR trafficking and cAMP signaling at endomembranes. J Cell Biol. 2025 Apr 7;224(4):e202409027.

Salmonowicz H, Szczepanowska K. The fate of mitochondrial respiratory complexes in aging. Trends Cell Biol. 2025 Mar 26:S0962-8924(25)00042-X.

Santos, R.; et al. A comprehensive map of molecular drug targets. Nat. Rev. Drug Discov 2017, 16, 19−34.

Scheidelaar S, Koorengevel MC, van Walree CA, Dominguez JJ, Dörr JM, Killian JA. Effect of Polymer Composition and pH on Membrane Solubilization by Styrene-Maleic Acid Copolymers. Biophys J. 2016 Nov 1;111(9):1974–1986. doi: 10.1016/j.bpj.2016.09.025.

Schöneberg J, Pavlin MR, Yan S, Righini M, Lee IH, Carlson LA, Bahrami AH, Goldman DH, Ren X, Hummer G, Bustamante C, Hurley JH. ATP-dependent force generation and membrane scission by ESCRT-III and Vps4. Science. 2018 Dec 21;362(6421):1423–1428.

Sligar SG, Denisov IG. Nanodiscs: A toolkit for membrane protein science. Protein Sci. 2021 Feb;30(2):297–315.

Souissi M, Pernier J, Rossier O, Giannone G, Le Clainche C, Helfer E, Sengupta K. Integrin-Functionalised Giant Unilamellar Vesicles via Gel-Assisted Formation: Good Practices and Pitfalls. Int J Mol Sci. 2021 Jun 13;22(12):6335. doi: 10.3390/ijms22126335.

Souza DP, Espadas J, Chaaban S, Moody ERR, Hatano T, Balasubramanian M, Williams TA, Roux A, Baum B. Asgard archaea reveal the conserved principles of ESCRT-III membrane remodeling. Sci Adv. 2025 Feb 7;11(6):eads5255.

Stephan T, Brüser C, Deckers M, Steyer AM, Balzarotti F, Barbot M, Behr TS, Heim G, Hübner W, Ilgen P, Lange F, Pacheu-Grau D, Pape JK, Stoldt S, Huser T, Hell SW, Möbius W, Rehling P, Riedel D, Jakobs S. MICOS assembly controls mitochondrial inner membrane remodeling and crista junction redistribution to mediate cristae formation. EMBO J. 2020 Jul 15;39(14):e104105.

Stephan T, Stoldt S, Barbot M, Carney TD, Lange F, Bates M, Bou Dib P, Inamdar K, Shcherbata HR, Meinecke M, Riedel D, Dennerlein S, Rehling P, Jakobs S. Drosophila MIC10b can polymerize into cristae-shaping filaments. Life Sci Alliance. 2024 Jan 22;7(4):e202302177.

Stępień P, Świątek S, Robles MYY, Markiewicz-Mizera J, Balakrishnan D, Inaba-Inoue S, De Vries AH, Beis K, Marrink SJ, Heddle JG. CRAFTing Delivery of Membrane Proteins into Protocells using Nanodiscs. ACS Appl Mater Interfaces. 2023 Nov 28;15(49):56689–701.

Strahl H, Ronneau S, González BS, Klutsch D, Schaffner-Barbero C, Hamoen LW. Transmembrane protein sorting driven by membrane curvature. Nat Commun. 2015 Nov 2;6:8728.

Streicher P, Nassoy P, Bärmann M, Dif A, Marchi-Artzner V, Brochard-Wyart F, Spatz J, Bassereau P. Integrin reconstituted in GUVs: a biomimetic system to study initial steps of cell spreading. Biochim Biophys Acta. 2009 Oct;1788(10):2291–300. doi: 10.1016/j.bbamem.2009.07.025.

Suzuki M, Kamiya K. Cell-sized asymmetric phospholipid-amphiphilic protein vesicles with growth, fission, and molecule transportation. iScience. 2023 Jan 31;26(3):106086. doi: 10.1016/j.isci.2023.106086.

Tang, B. et al. (2026). Structure and stability of styrene-maleic acid lipid particles: the role of polymer-lipid ratio. Journal of Colloid and Interface Science 724, Part 1, 141054. DOI: 10.1016/j.jcis.2026.141054

Tosaka T, Kamiya K. Formation of Cell-Sized Liposomes Incorporating a β-Barrel-Structured Porin through Rehydration of a Phospholipid-Membrane Protein Dried Film. ACS Omega. 2024 Jan 26;9(5):5911–5918. doi: 10.1021/acsomega.3c09431.

Van de Cauter L, Fanalista F, van Buren L, De Franceschi N, Godino E, Bouw S, Danelon C, Dekker C, Koenderink GH, Ganzinger KA. Optimized cDICE for Efficient Reconstitution of Biological Systems in Giant Unilamellar Vesicles. ACS Synth Biol. 2021 Jul 16;10(7):1690–1702.

Varnier A, Kermarrec F, Blesneac I, Moreau C, Liguori L, Lenormand JL, Picollet-D’hahan N. A simple method for the reconstitution of membrane proteins into giant unilamellar vesicles. J Membr Biol. 2010 Feb;233(1-3):85–92. doi: 10.1007/s00232-010-9227-8.

Yanagisawa M, Iwamoto M, Kato A, Yoshikawa K, Oiki S. Oriented reconstitution of a membrane protein in a giant unilamellar vesicle: experimental verification with the potassium channel KcsA. J Am Chem Soc. 2011 Aug 3;133(30):11774–9. doi: 10.1021/ja2040859.

