## Supplementary Figures 1 to 3 for "Reconstitution of Integral Membrane Proteins at Catenoid Necks via the Synthetic Membrane Transfer"

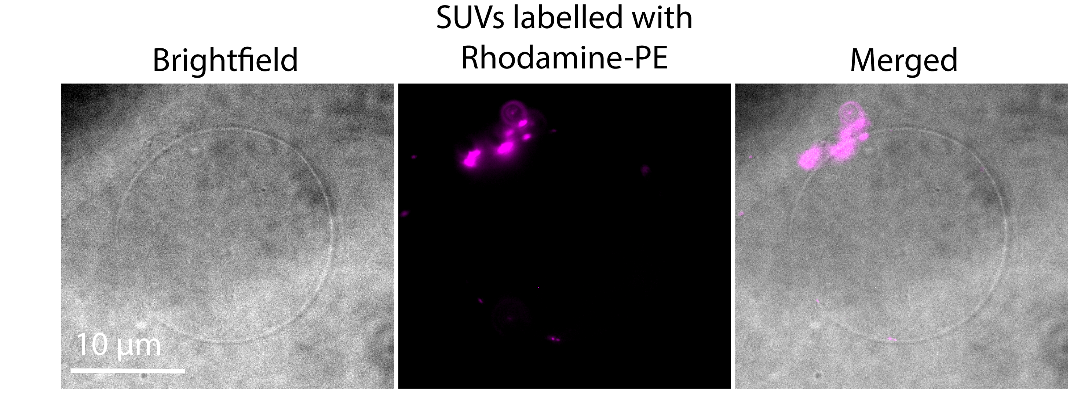

***Supplementary Figure 1.*** *Representative images of GUVs generated in the presence of DOPE–Rhodamine-labeled SUVs. SUVs fail to integrate into the membrane and instead form aggregates at the membrane, indicating inefficient lipid transfer compared to lipid-SMA.*

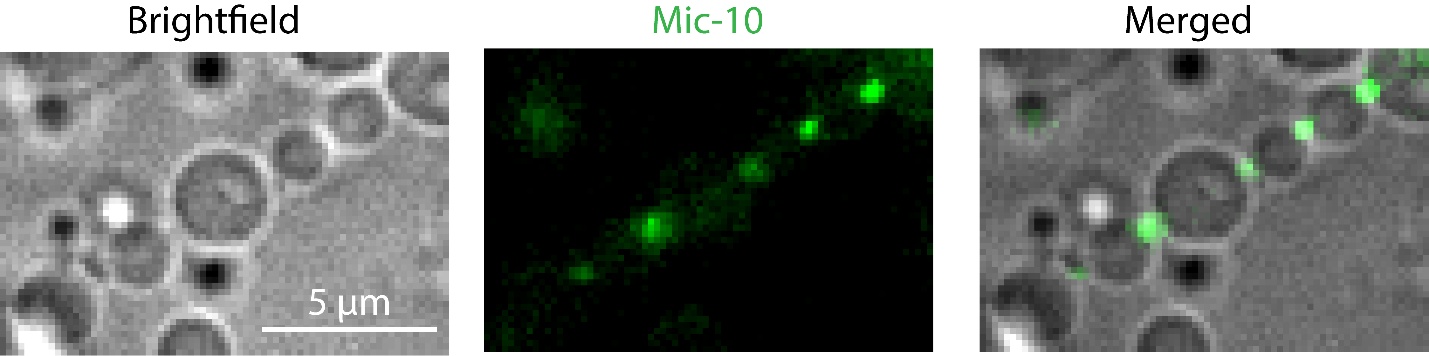

***Supplementary Figure 2.*** *Additional example of chain of dumbbell-shaped GUVs with reconstituted Mic10–SMA. Mic10 clusters are enriched at membrane necks connecting adjacent compartments.*

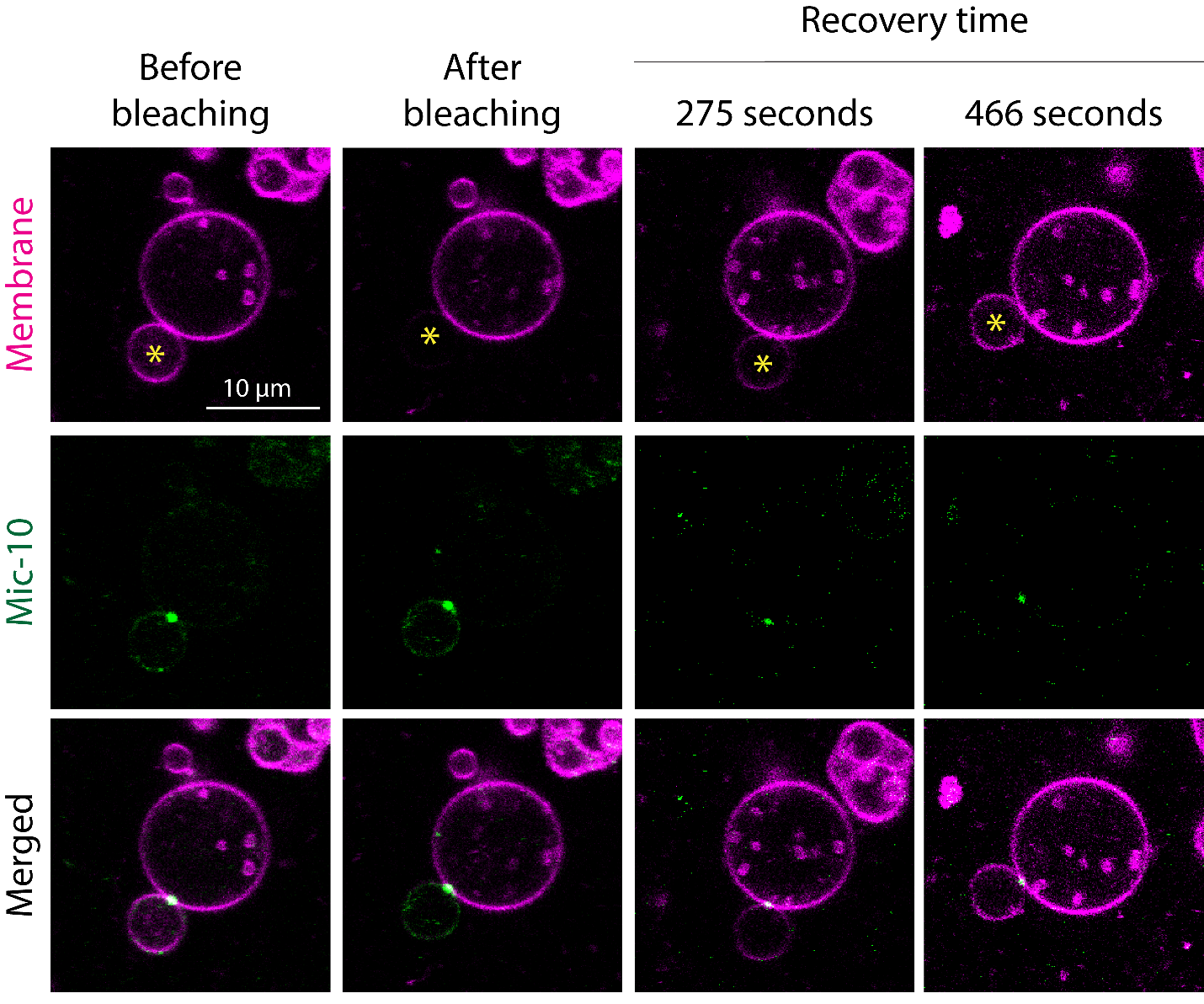

***Supplementary Figure 3.*** *FRAP analysis of a dumbbell GUV containing a reconstituted Mic10 cluster at the neck. Recovery of membrane fluorescence in the bleached lobe demonstrates membrane continuity through the neck.*

**Movie captions**

**Movie 1.** Time-lapse epifluorescence imaging of a spherical GUV with reconstituted GFP–Mic10. Mic10 forms dynamic clusters that diffuse along the membrane. GFP fluorescence is shown in green. Frame rate: 0.33 fps.

**Movie 2.** Time-lapse epifluorescence imaging of a GUV reshaped into a chain of dumbbells. Mic10 clusters are stably localized at membrane necks. Merged brightfield and GFP fluorescence (green). Frame rate: 0.33 fps.

**Movie 3.** Time-lapse epifluorescence imaging of an independent example of a dumbbell-shaped GUV. Mic10 clusters remain enriched at membrane necks. Merged brightfield and GFP fluorescence (green). Frame rate: 0.33 fps.

**Movie 4.** Time-lapse confocal imaging of nanotubes emanating from a GUV. Mic10 clusters localize at the nanotube neck. Merged lipid fluorescence (magenta) and GFP fluorescence (green). Frame rate: 1 fps.
